# CRISPR-Cas interference decays rapidly with distance from the leader sequence in a long array

**DOI:** 10.64898/2026.07.09.737519

**Authors:** Marijn H. Ceelen, Mauro Albertini, Gregory J. Velicer, Sébastien Wielgoss

## Abstract

Spacer efficacy generally declines with distance from the leader sequence, but the scarcity of fine-scale studies hampers comparisons across taxa. Here, we investigated positional effects across an exceptionally long 121-spacer CRISPR array associated with the type I-C *cas* operon of a *Myxococcus xanthus* natural isolate. In plasmid-interference assays, we found that interference rapidly declined with distance from the leader sequence, with only the proximal ∼4% of spacers conferring measurable interference. This contrasts strikingly with a study in *Vibrio cholerae*, in which it was shown that ∼95% of spacers in a shorter (39-spacer) array were effective. Our results suggest that there is great variation in the effective proportion of spacers across species, highlighting the need for fine-scale studies of CRISPR-array activity across diverse bacterial lineages.

## Introduction

Bacteria frequently encounter mobile genetic elements (MGEs), which can be beneficial (*e*.*g*., plasmids with antibiotic resistance genes), deleterious (*e*.*g*., costly plasmids with toxin-antitoxin systems) or lethal (*e*.*g*., lytic bacteriophage). In response to harmful MGEs, many defense systems have evolved.^1^ These include CRISPR-Cas defense systems, in which a CRISPR array stores short sequences from previously encountered foreign DNA − known as spacers − between identical direct repeat sequences. Spacers derive from protospacers that are flanked by a characteristic protospacer adjacent motif (PAM), a crucial marker that signifies a protospacer sequence as “non-self”.^2,3^ In a process known as interference, CRISPR arrays are transcribed and processed into crRNAs that guide Cas proteins to complementary target phage DNA sequences, which are then cleaved, preventing further replication. While these defense platforms are highly diverse and categorized into distinct classes and types based on their effector module composition, they universally rely on the structural integrity of the transcribed array.^4^

New spacers are usually integrated into the CRISPR array near the leader sequence at the 5′ end (called proximal), which contains the promoter.^5^ We expect that newer spacers are more beneficial on average, as they likely often derive from more recently encountered phage. In contrast, older spacers may be less likely to be effective against currently circulating invasive genetic elements, and might thus represent unnecessary genetic cargo. This could explain the observation that older spacers are regularly pruned by deletions and reshuffled by recombination events.^6^ Additionally, spacers close to the 5′ end generally are more effective against invading DNA than spacers very distant from the leader sequence (called distal).^7–9^

This difference in efficacy may be explained by the higher net expression of proximal spacers than of distal spacers.^7,8,10–12^ Given that the entire CRISPR array is transcribed as a single unit, early termination events may reduce the overall expression of spacers further downstream. As CRISPR-array transcripts are non-coding and can be very long, they have been proposed to be particularly susceptible to Rho-dependent transcription termination.^13,14^ In many bacterial systems, premature Rho termination can be counteracted by a *boxA*-dependent Nus factor complex.^15^ *boxA* sequences are also commonly found in CRISPR leader regions, where they facilitate antitermination and promote complete transcription of CRISPR arrays. Nevertheless, premature Rho termination has been shown to impair the utilization of distal spacers and reduce CRISPR interference even in the presence of *boxA*-mediated antitermination.^14^ Crucially, however, the sequence context of these motifs matters; in one case in *Salmonella*, the recruitment of Nus-factors via *boxA* enhanced Rho-dependent termination instead of preventing it.^16^

Additionally, earlier-transcribed spacer-repeat units are more likely to be processed first and would have priority access to unbound effector protein complexes to initiate spacer-guided DNA interference.^17^ In line with these expectations, McGinn and Marraffini demonstrated that spacer integration in the middle of a CRISPR array provided less resistance than integration at the start of the array.^8^ Similarly, Rao et al. reported an ∼100-fold increase in CRISPR interference efficacy against a matching protospacer by moving the targeting spacer (Sp1) from the most distal to the most proximal position by deleting the 43 preceding spacer-repeat tandems, leaving only Sp1.^9^

Despite the more general observations outlined above, fine-scale effects of spacer position remain largely unexplored. The work by Stringer *et al*. (2020) stands out for testing the interference activity of each spacer in a 39-spacer array in a *Vibrio cholerae* strain with an intact *boxA* sequence. They found that all but the two most distal spacers mediated high levels of interference.^14^ This result contrasts with studies indicating that mid-array spacers can have reduced efficacy,^6,8^ suggesting that the relationship between spacer position and efficacy may be highly variable.

The number of spacers in CRISPR arrays varies greatly, from just one to hundreds.^18^ The median number of spacers in type I CRISPR arrays has been reported to be 26,^19^ which is similar to that for CRISPR-Cas systems in bacteria generally.^18^ Arrays in the majority of genomes with a CRISPR-Cas system have 50 or fewer spacers (∼74%), but ∼8% of genomes with CRISPR-Cas have arrays with more than 100.^18^ Whether and how the relationship between spacer position and efficacy covaries with array length is unknown. If spacer efficacy were to decrease similarly as a function of absolute spacer position across short and long arrays, this would increase the puzzle of why many genomes have very long arrays. If the relationship were to differ fundamentally between long vs short arrays, this would raise the intriguing question of how.

A compelling model species for increasing our understanding of spacer-position effects is *Myxococcus xanthus*, a soil bacterium renowned for its complex social traits, including cooperative predation and multicellular fruiting-body development. *M. xanthus* genomes typically carry multiple CRISPR-Cas systems with exceptionally long CRISPR arrays.^20,21^ CRISPR-Cas systems are categorized into two classes based on the composition of the effector complex by multiple proteins (class 1) or a single protein (class 2). These classes are further divided into multiple types, with, among others, type I (targeting DNA) and type III (targeting RNA) belonging to class 1 and type II (targeting DNA) belonging to class 2.^4^ Three such systems are encoded in parallel in common lab strains of *M. xanthus*, a type I-A system linked to the regulation of development,^21,22^ a highly conserved type I-C system which was demonstrated to be actively involved in defense,^23^ and a type-III system that acts in conjunction with a neighboring CBASS system.^24^ Crucially, natural isolates of *M. xanthus* obtained from the same local soil population exhibit striking variation in their younger type I-C spacers, indicating recent and distinctive ecological interactions with local mobile genetic elements (MGEs).^20^

Here, we set out to characterize the relationship between spacer position and interference activity in a relatively long array at a fine positional scale. To do this, we utilized the *M. xanthus* strain GH3.5.6c02, a natural isolate for which extensive phenotypic and genomic data exist.^20,25^ Unlike common laboratory strains (such as DK1622), which maintain multiple overlapping systems, GH3.5.6c02 contains only a single type I-C *cas* operon associated with an expanded array of 121 spacers.^20^ This strain is one of many natural isolates that were collected to enable investigation of genetic and phenotypic variation among *M. xanthus* strains across a range of geographic scales.^20,26–29^

We systematically mapped the interference efficacy across its extensive CRISPR array, with a high-density sampling strategy focused on the leader-proximal region (spacers 121 to 105), extending our assay just past the threshold where interference was not measurably different from a negative-control strain with debilitated CRISPR-Cas I-C function. Investigating the effect of position on interference efficacy in this way will improve our general understanding of spacer-position effects and help with the interpretation of the biological importance and evolutionary utility of the many spacers stored in long CRISPR arrays.

## Materials & Methods

### Construction of a type I-C CRISPR knockout (KO) strain

A 524bp fragment of the GH3.5.6c02 I-C *cas3* gene was amplified by PCR using primers cas3_For and cas3_Rev. (For a full list of strains (supplementary table 1), plasmids (supplementary table 2), and primers/oligonucleotides (supplementary table 3) used in this paper, refer to the supplementary material.) This fragment was then ligated into plasmid pCR-Blunt (blunt-ended ligation) and transformed by heat shock into *Escherichia coli* One Shot Top10 chemically competent cells (Invitrogen). The transformed *E. coli* cells were allowed to recover in 1 mL LB medium for 1 hour at 37 °C and spread on LB agar (1.5%) kanamycin (40 µg/mL) plates. Correct insertion was confirmed by running EcoRI-digested plasmid on a 1% agarose gel and observing the expected restriction pattern (two bands of 524 bp and 3.5 kb) and by Sanger sequencing (using standard primer M13R on the vector).

Competent cells of *M. xanthus* GH3.5.6c02 were prepared as follows: 8 mL of fresh cell cultures were collected at mid-exponential growth phase (OD 0.2-0.8) and centrifuged for 15 minutes at 4500g (Eppendorf 5810R centrifuge). After discarding the supernatant, the cell pellet was resuspended in 30 mL of demineralized water. After another round of centrifugation and discarding the supernatant, the pellet was resuspended in 2 mL of demineralized water. This step was followed by three additional washing steps comprising centrifugation for 3 minutes at 20,800g (Eppendorf 5417C centrifuge), the discarding of supernatants, and resuspension in 2 mL of demineralized water. After the final step, the pellet was spun down and resuspended in 200 µL of demineralized water. For each transformation, 3 µL of plasmid solution was added to 60 µL of competent cells. Cells were then transferred to a standard electroporation cuvette (gap width 1 mm) and electroporated at 400 Ω, 25 µFd, and 0.65 kV (BioRad Genepulser II). The electroporated cells were then allowed to recover overnight at 32°C, 300 rpm in 2.5 mL CTT-liquid. Recovered cells were grown for 3 more days in 8 mL CTT-liquid containing kanamycin (40 µg/mL), and diluted on CTT-agar (0.5%) plates containing kanamycin (40 µg/mL) incubated at 32°C for 5-7 days. Single colonies were picked, grown in CTT liquid and stored in 20% (v/v) glycerol stocks at -70°C. Clones were subsequently tested for developmental and swarming capability relative to the original WT strain. A single I-C-KO clone that did not show deficiencies at development or swarming was chosen for the plasmid interference assays in this paper and named GH3.5.6c02_*cas3*::pCR-Blunt.

### Cloning of protospacer plasmids

The tetracycline resistance gene on plasmid pMR3629 ^30^ was amplified with primers GV1002 and GV1003 using Phusion polymerase (ThermoFisher). The PCR product and pZJY41 vector^31,32^ were then both digested with XbaI and BamHI (NEB) and ligated with T4 ligase (NEB) to construct plasmid pMxtetR. The resulting plasmid was used to transform *E. coli* One Shot Top10 chemically competent cells (Invitrogen) by heat shock.

Protospacer sequences with type-I-C-consensus PAM-motif (CTTC; Hu et al. 2024) were ordered as ssDNA oligos (Microsynth) and annealed by heating to 95°C and cooling down to 25°C for one hour to form dsDNA molecules with overhangs. The annealed oligos were then ligated into BamHI+KpnI (NEB) digested plasmid pMxtetR, which was then used to transform *E. coli* One Shot Top10 chemically competent cells (Invitrogen).

### Culturing conditions

*E. coli* was grown in LB medium at 32°C and 300 rpm in liquid and on LB agar (1.5%) plates at 32°C and 90% rH. *M. xanthus* was grown in CTT medium at 32°C and 300 rpm and on CTT-agar plates at 32°C and 90% rH.

### Plasmid Interference Assay

To test the relative transformation efficiency of the different plasmids carrying a protospacer matching the CRISPR arrays of *M. xanthus* in the WT vs I-C-KO backgrounds, *M. xanthus* cells were grown overnight in liquid CTT medium to the mid-log growth phase. Optical density was then measured at 595 nm, and the number of cells/mL was estimated. An estimated 8*10^8^ cells (∼2 mL culture at OD 0.45) were used per transformation. The cells were then centrifuged for 5 minutes at 4472g. The supernatant was discarded and the cell pellet was resuspended in 2 mL of demineralized water. The resuspended cells were then centrifuged for 5 minutes at 4472g again. This was repeated for a total of four washing steps. The cells were then resuspended in 80 µL of demineralized water. For each transformation, 100 pmol of plasmid was added and mixed with the cells by tapping. In the no-plasmid treatment, 2.5 µL of water was added instead. The cells were then pipetted into a 1 mm electroporation cuvette (BTX, Cambridge, UK) and electroporated at 400 Ω, 25 µF, 0.65 kV. A volume of 920 µL of CTT medium was added to the cuvette, and 1 mL was transferred to a flask containing 1.5 mL of CTT medium. The cells were then allowed to recover for 4 hours at 32°C and 300 rpm. After recovery, the cells were serially diluted in CTT medium, and 100 µL was mixed with 10 mL of CTT-SA (0.5 % agar) with or without antibiotic (12.5 µg/mL oxytetracycline). After cooling to solidify for 30 minutes, the plates were incubated at 32°C and 90% rH for 4 or 5 days, followed by colony counting. The concentration of cells as CFU (Colony Forming Units)/mL was calculated from the plated volume and the dilution factor. The transformation efficiency was then calculated by the formula: *transformation efficiency* 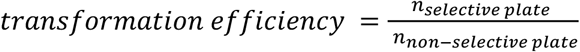, where *n* is the calculated CFU/mL.

### Statistics

To analyze the effect of strain and position on log_10_-transformed transformation efficiency, we fit a linear mixed-effects model (LMEM) using the lme4 package (version 1.1-37) in R (version 4.6.0).^33^ The model included strain, position, and their interaction as fixed effects, and replicate date as a random effect to account for batch variation. After fitting the LMEM, pairwise comparisons of WT vs. KO strains at each position were performed using estimated marginal means (EMMs) from the emmeans R package (version 1.11.2-8) with a Holm-Bonferroni adjustment for multiple comparisons.^34^

## Results

To measure the degree of CRISPR interference, we constructed an isogenic control strain, the null-expression-knockout mutant GH3.5.6c02_*cas3*::pCR-Blunt, hereafter referred to as I-C-KO, in which interference has been debilitated by disruption of the *cas3* gene. We then measured the transformation efficiency of plasmids carrying a protospacer matching the CRISPR array with accompanying PAM, in both the WT and the I-C-KO strains, and scored spacer efficacy, which we define as the ability of a spacer to reduce plasmid transformation efficiency.^35^ We tested a fine-scale range of spacer positions at the promoter-proximal end of the array (Figure 1), starting with the most proximal spacer 121. Transformation efficiency was calculated from the number of colonies that grew with vs. without a selective antibiotic. As controls, we used the empty vector and a plasmid carrying a non-matching protospacer. We chose GH3.5.6c02 as a wild-type (WT) strain because it carries only one CRISPR-Cas system, whereas standard lab strains (such as DK1622) and most natural isolates sequenced so far have multiple systems.^20^ Hence, we removed the possibility of crosstalk between CRISPR-Cas systems that might confound measurements of CRISPR-Cas interference.

**Figure 1.**
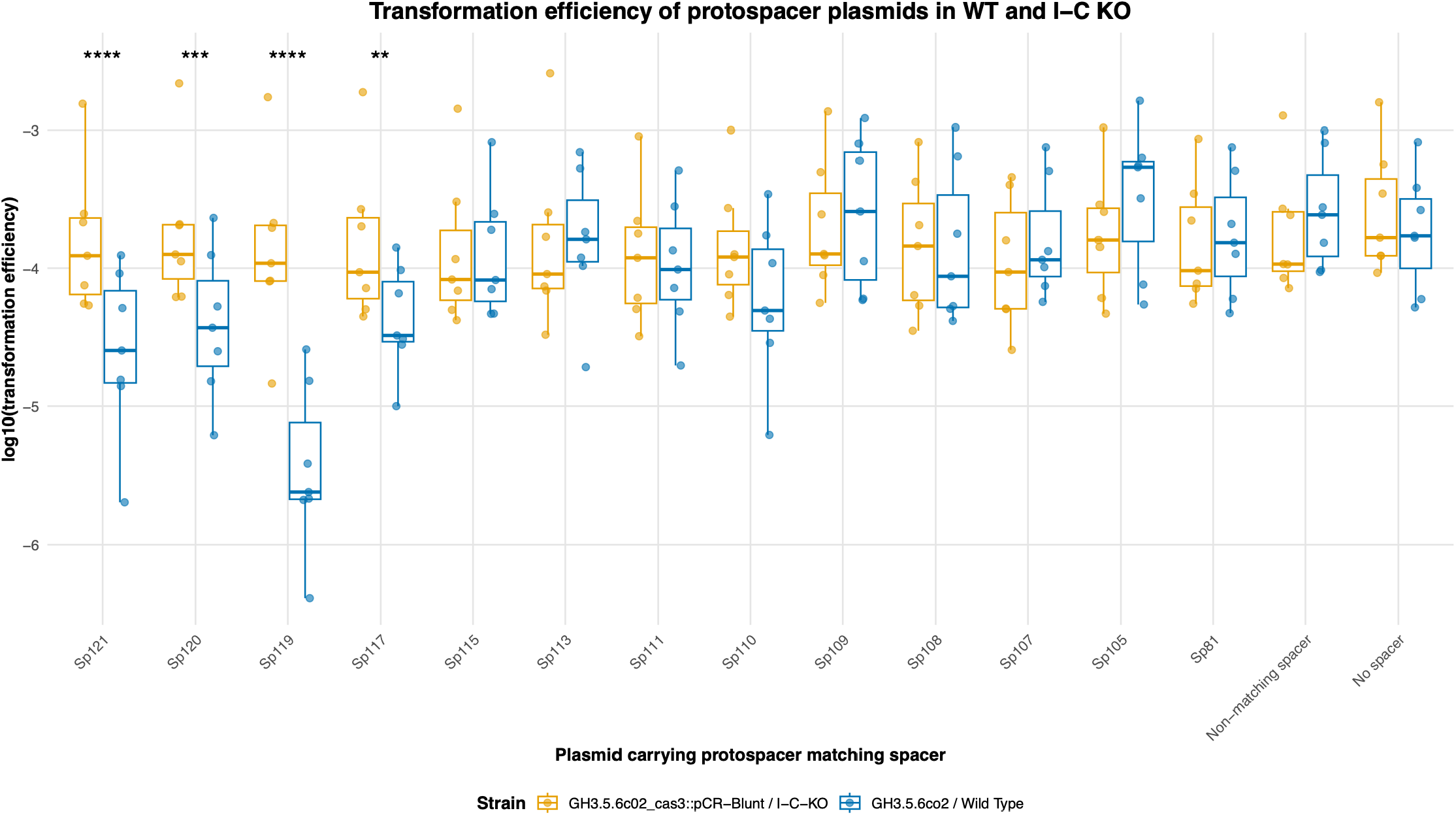
Transformation efficiencies of plasmids carrying different protospacers into WT (strain GH3.5.6c02) and a I-C-KO strain (GH3.5.6c02_cas3::pCR-Blunt). Spacer numbers indicate position within the array, counting down from the leader sequence. In the WT background, transformation efficiency varies greatly between plasmids with different spacers, but such variation is not observed in the I-C-KO background. Asterisks indicate significant differences between the transformation efficiency of a given plasmid in WT vs I-C-KO (**** = p < 0.0001, *** = p < 0.001, and ** = p < 0.01). The non-matching spacer corresponds to Spacer 1 in the I-C CRISPR array of strain M. xanthus DK1622 and does not match any spacer in the GH3.5.6c02 CRISPR array. The no-spacer treatment corresponds to the empty vector pMxtetR.

While plasmid transformation efficiencies varied little across tested spacers in the CRISPR-knockout strain (figure 1), they varied strongly in the WT strain. This demonstrates that the type-I-C CRISPR-Cas system is active in the natural isolate under laboratory conditions and that the system’s activity was disrupted in the I-C-KO strain. Using a linear mixed-effects model (LMEM) and contrasting the estimated log-transformed transformation efficiency in the I-C-KO and WT strains (emmeans, Holm-Bonferroni correction), we find that the transformation efficiency is significantly lower in the WT background for the four most proximal tested spacers: 121, 120, 119, and 117. Importantly, none of the spacers beyond position 117 show a significantly different transformation efficiency between the WT and I-C-KO backgrounds after correcting for multiple testing (*p*_corrected_ > 0.05 each, figure 1). We then plotted the relative transformation efficiency for each spacer, calculated by dividing the transformation efficiency for each spacer plasmid in the WT strain by that in the I-C-KO strain, against the distance of the spacer from the leader sequence (figure 2). As illustrated in figure 2, spacer efficacy decays rapidly between five and seven spacers from the leader sequence. Additionally, the relative transformation efficiency varies strongly among the effective spacers. Spacer 119 stands out for its substantially lower average relative transformation efficiency of 5*10^-2^, in contrast to spacers 121, 120, and 117, which have average relative transformation efficiencies of 3*10^-1^, 3*10^-1^, and 5*10^-1^ respectively.

**Figure 2.**
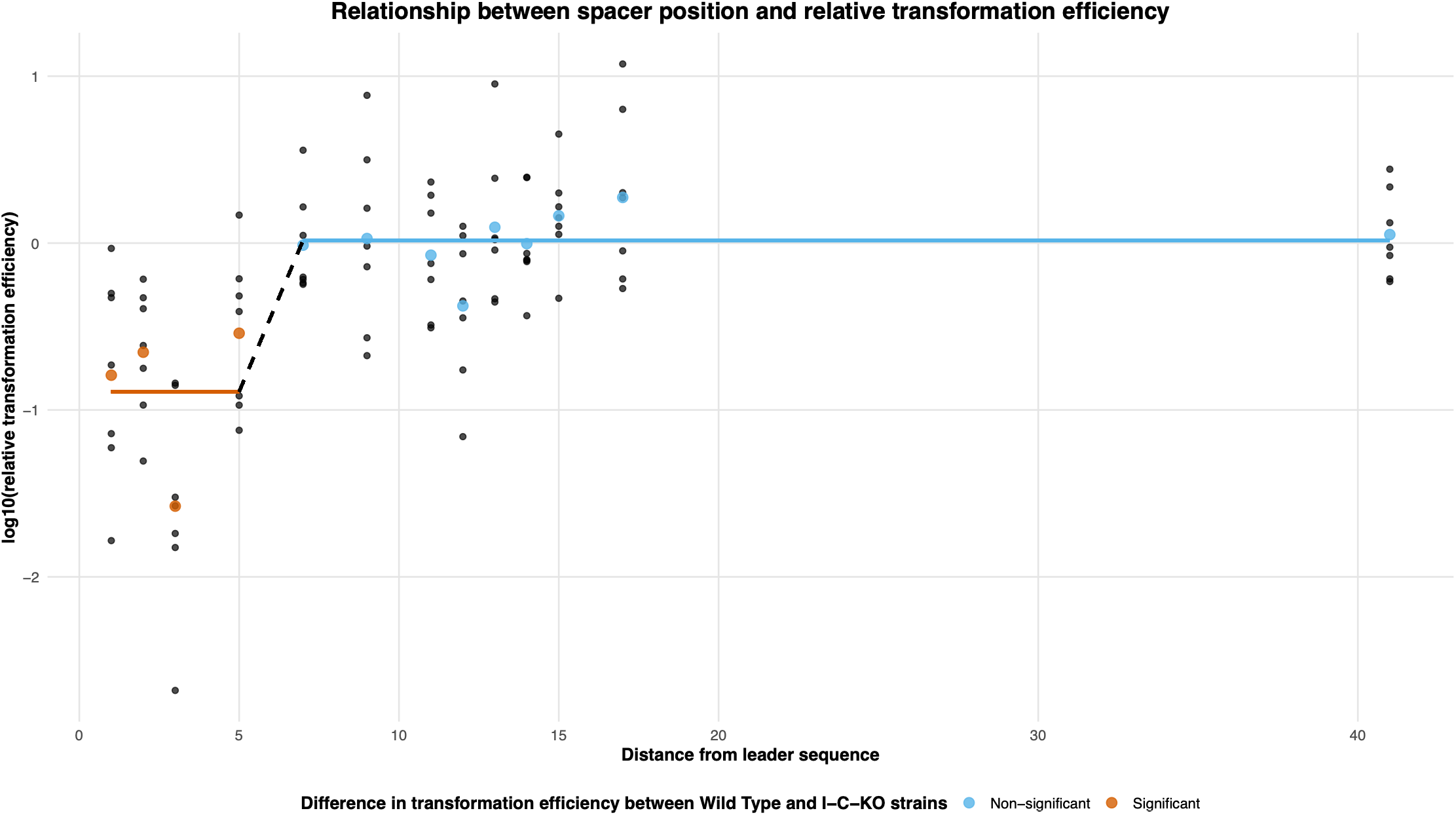
Relative transformation efficiency of plasmids carrying protospacers as a function of the distance of the matching spacer from the leader sequence (counted by the number of repeat sequences between spacer and leader sequence). Relative transformation efficiency was calculated by dividing the transformation efficiency in the WT strain by that in the I-C-KO strain. A log10-transformed relative transformation efficiency of zero means that there is no difference in transformation efficiency between the two strains. The red and blue points represent the average value for each spacer plasmid and the red and blue lines represent the average for spacers with a significant and non-significant difference between the WT and I-C-KO strain, as calculated previously, respectively.

## Discussion

Here we have characterized how spacer-guided CRISPR interference efficacy decreases with distance from the leader sequence at very fine scales in *Myxococcus xanthus*, notorious for its exceptionally long CRISPR spacer arrays. Only the spacers within the five most proximal positions (positions 117-121) showed strong interference, whereas tested spacers from position 115 onwards showed no significant interference. These observations are consistent with previous coarser-grained studies of positional effects^7,8,11,12^ but contrast strikingly with a fine-scale investigation of all 39 spacers in a CRISPR array associated with a type I-E *cas* operon in *Vibrio cholerae*. In the latter study, the 35 most promoter-proximal spacers showed activity while only the oldest two did not.^14^ While we cannot exclude that distal spacers may show higher effectiveness under natural conditions, in comparison to previous studies our results suggest that there may be large variation between strains and arrays in the effect of distance on spacer efficacy. In some cases, large fractions of CRISPR arrays may be ineffective at interference.

*boxA*-mediated antitermination plays a crucial role in promoting the complete transcription of CRISPR arrays in genera like *Salmonella* and *Vibrio*.^14^ Importantly, a highly conserved *boxA* sequence is also present in the leader sequence of the type I-C CRISPR-Cas system in *M. xanthus* strain GH3.5.6c02 (*M. xanthus* motif: GCTCTTTGAAA; *E. coli* consensus: GNTCTTTAANA )^14^. Furthermore, this exact sequence is identical to the *boxA* motif found upstream of the *M. xanthus* 16S rRNA (*rrnA*) gene, and homologs of the complete Nus-factor complex are fully encoded within the strain’s genome. Conventionally, the presence of these components would imply robust antitermination, suggesting that the observed low activity of downstream distal spacers cannot be attributed to structural termination. Intriguingly, however, evidence from *Salmonella* demonstrates that the *boxA*-NusB interaction is not strictly anti-terminating; under certain genetic architectures, NusB recruitment via *boxA* can actively enhance Rho-dependent termination at sub-optimal rut sites, as seen in the *hisG* cistron.^16^ If this is the case here, it raises the interesting question as to why transcription of the array would be terminated early.

In our results, spacer activity does not decrease uniformly with distance from the leader sequence. For example, spacer 119 apparently exerts more interference than the more proximal spacers 121 and 120 (figure 1). Such non-linearity might be explained by spacer-sequence effects, as spacer sequence has been shown to affect the formation of secondary structures that are important for the binding of crRNA to the Cas proteins.^36,37^ Additionally, the GC-content of the spacer may play a role, with Xu Hua Fu et al. finding an optimum at ∼62.5% in type I-E.^38^

Our finding that only a small portion of a long *M. xanthus* CRISPR array shows evidence of being active in interference raises the question of why long arrays persist in bacterial populations. Due to large effective population sizes, selection among bacteria has been estimated to be efficient enough to remove insertions as small as 10 bp due to their added replication costs,^39^ consistent with the hypothesis that long CRISPR arrays are maintained by selection. Selection on genome size may be somewhat relaxed in *M. xanthus*, which has a large genome (∼9 Mb) combined with a relatively small effective population size at ∼10^7^.^40^ However, there are species with much smaller genomes and longer arrays (*e*.*g*., *Sulfolobus tokodaii*, 2.6 Mb genome, 458 spacers).^41^ If non-interfering spacers are maintained by selection, this raises the question of how they are beneficial. Two mechanisms that might mediate the benefits of non-interfering spacers are recombination and primed adaptation (priming).

Firstly, genomic rearrangement, for example by recombination, may move non-interfering spacers to the proximal end of the array and thereby activate them.^9^ Additionally, horizontal gene transfer (HGT) may allow divergent strains to share a large, population-level spacer library, allowing them to respond more effectively to rare threats. The repeating structure of CRISPR arrays facilitates recombination events. As an example, ∼20% of spacers from an *E. coli* strain isolated from a frozen mammoth calf matched modern *E. coli* spacers. However, these spacers were not concentrated at the distal end of the array and were instead scattered through the array, suggesting reshuffling of spacers.^42^ *M. xanthus* is known to undergo a high degree of HGT, which has also been shown to occur within the CRISPR-Cas operons.^43–45^

Secondly, ineffective spacers can increase uptake of new spacers through priming. Priming can occur when a CRISPR ribonucleoprotein (crRNP) effector complex recognizes a protospacer target, which then increases spacer uptake from nearby DNA.^46^ Strong interference is not required for priming, as mismatched spacers and non-canonical PAMs can support priming, which Fineran et al. suggested may support the retention of spacers that have become obsolete through mutation.^47^ Deecker et al. showed that distal spacers with low interference can assist spacer acquisition.^7^ We suggest that priming could thus help explain the maintenance of long CRISPR arrays. Recombination and priming can allow bacteria to use long arrays to adapt faster to new threats. In this way, inactive spacers may contribute to the bacterial immune system.

In addition to raising the question of why long arrays persist, our findings may help explain two previously puzzling results. Firstly, self-targeting spacers, which target the bacterial genome and result in cell death when effective, were found to be widespread; Despite this, CRISPR-Cas systems are generally maintained.^48^ Self-targeting spacers that are located in inactive sections of CRISPR arrays may be tolerable, partially explaining this finding. Secondly, while CRISPR-Cas systems have been theorized to form a barrier to HGT, Gophna et al. found no correlation between the number of spacers and the number of genes suspected of HGT.^49^ Our findings suggest that spacer count is a poor proxy for CRISPR-Cas activity, which may help to explain the findings by Gophna *et al*.

## Conclusions

To conclude, this paper demonstrates that only a very limited number of proximal CRISPR spacers tested in a long, 121-spacer array are effective at generating interference of a DNA target within the native genetic background of an *M. xanthus* natural isolate. Given the large differences in the effect of distance from the leader sequence on spacer efficacy observed across our study, several coarse-grained studies, and one other fine-scaled study,^7,8,11,12,14^ more high-resolution studies are critically needed to map what fractions of spacers provide meaningful resistance to MGEs across diverse microbial lineages. In particular, future research must look beyond the mere presence of regulatory motifs to scrutinize their exact sequence context. It will be of interest to investigate how similar or variable the relationship between distance from the leader sequence and efficacy is across bacterial strains and species, and between short vs long arrays, as well as the evolutionary mechanisms by which ineffective portions of longer CRISPR arrays persist.

## Acknowledgments

The authors thank Lasse Nielsen and Dr. Yuen-Tsu Nicco Yu for creating the GH3.5.6c02_*cas3*::pCR-Blunt strain and Beatriz Menezes de Albuquerque for collecting preliminary data.

## Author contributions

### Statements and declarations

#### Ethical considerations

ethical approval was not required for this study.

Consent to participate: Not applicable.

Consent for publication: Not applicable.

#### Author Disclosure Statement

The author(s) declared no potential conflicts of interest with respect to the research, authorship, and/or publication of this article.

#### Funding

This research was funded by the Swiss National Science Foundation (grant 310030_207923).

## Notes

### Competing Interest Statement

The authors have declared no competing interest.

