## Supplemental Table 1, 2, and 3 for "CRISPR-Cas interference decays rapidly with distance from the leader sequence in a long array"

### Supplementary data

##### *Table 1. Strains*

| No. | Species | Strain | Use | Source | Notes |
|---|---|---|---|---|---|
| **1** | *Escherichia coli* | Top10 | Cloning | Invitrogen | |
| **2** | *Myxococcus xanthus* | GH3.5.6c02 | Plasmid interference assay | Natural isolate (Kraemer & Velicer, 2011, PNAS) | |
| **3** | *Myxococcus xanthus* | GH3.5.6c02_*cas3*::pCR-Blunt | Plasmid interference assay | Genetically modified natural isolate | CRISPR-Cas disabled by insertion of pCR-Blunt |

##### *Table 2. Plasmids*

| No. | Name | Source | Use | Origin of replication | Size (bp) | Antibiotic resistance |
|---|---|---|---|---|---|---|
| **1** | pCR-Blunt | ThermoFisher Scientific | Cloning vector | ColE1 | 3512 | Kanamycin, Bleomycin |
| **2** | pCR-Blunt_Cas3 | Modified pCR-Blunt | GH3.5.6c02 Cas3 insertion | ColE1 | 4036 | Kanamycin, Bleomycin |
| **3** | pZJY41 | State Key Laboratory of Microbial Technology, Shandong University, China (Feng et al., 2012; Zhao et al., 2008) | Cloning vector | ColE1 (*E. coli*) and pMF1 (*M. xanthus*) | 6740 | Ampicillin, Kanamycin |
| **4** | pMR3629 | Departamento de Genética y Microbiología, Área de Genética, Universidad de Murcia, Spain (Iniesta et al., 2012) | Source tetR gene | X | X | X |
| **5** | pMxTetr | Modified pZJY41 by addition of Oxytetracycline resistance | Cloning vector and plasmid interference control | ColE1 (*E. coli*) and pMF1 (*M. xanthus*) | 8723 | Oxytetracycline, Ampicillin, Kanamycin |
| **6** | pMxtetR_Sp121 | pMxtetR with GH3.5.6c02 protospacer 121 inserted in BamHI+KpnI site | Plasmid interference assay | ColE1 (*E. coli*) and pMF1 (*M. xanthus*) | 8756 | Oxytetracycline, Ampicillin, Kanamycin |
| **7** | pMxtetR_Sp120 | pMxtetR with GH3.5.6c02 protospacer 120 inserted in BamHI+KpnI site | Plasmid interference assay | ColE1 (*E. coli*) and pMF1 (*M. xanthus*) | 8755 | Oxytetracycline, Ampicillin, Kanamycin |
| **8** | pMxtetR_Sp119 | pMxtetR with GH3.5.6c02 protospacer 119 | Plasmid interference assay | ColE1 (*E. coli*) and | 8756 | Oxytetracycline, Ampicillin, Kanamycin |

| | | | | | | |
|---|---|---|---|---|---|---|
| | | inserted in BamHI+KpnI site | | pMF1 (*M. xanthus*) | | |
| **9** | pMxtetR_Sp117 | pMxtetR with GH3.5.6c02 protospacer 117 inserted in BamHI+KpnI site | Plasmid interference assay | ColE1 (*E. coli*) and pMF1 (*M. xanthus*) | 8759 | Oxytetracycline, Ampicillin, Kanamycin |
| **10** | pMxtetR_Sp115 | pMxtetR with GH3.5.6c02 protospacer 115 inserted in BamHI+KpnI site | Plasmid interference assay | ColE1 (*E. coli*) and pMF1 (*M. xanthus*) | 8757 | Oxytetracycline, Ampicillin, Kanamycin |
| **11** | pMxtetR_Sp113 | pMxtetR with GH3.5.6c02 protospacer 113 inserted in BamHI+KpnI site | Plasmid interference assay | ColE1 (*E. coli*) and pMF1 (*M. xanthus*) | 8757 | Oxytetracycline, Ampicillin, Kanamycin |
| **12** | pMxtetR_Sp111 | pMxtetR with GH3.5.6c02 protospacer 111 inserted in BamHI+KpnI site | Plasmid interference assay | ColE1 (*E. coli*) and pMF1 (*M. xanthus*) | 8755 | Oxytetracycline, Ampicillin, Kanamycin |
| **13** | pMxtetR_Sp110 | pMxtetR with GH3.5.6c02 protospacer 110 inserted in BamHI+KpnI site | Plasmid interference assay | ColE1 (*E. coli*) and pMF1 (*M. xanthus*) | 8757 | Oxytetracycline, Ampicillin, Kanamycin |
| **14** | pMxtetR_Sp109 | pMxtetR with GH3.5.6c02 protospacer 109 inserted in BamHI+KpnI site | Plasmid interference assay | ColE1 (*E. coli*) and pMF1 (*M. xanthus*) | 8758 | Oxytetracycline, Ampicillin, Kanamycin |
| **15** | pMxtetR_Sp108 | pMxtetR with GH3.5.6c02 protospacer 108 inserted in BamHI+KpnI site | Plasmid interference assay | ColE1 (*E. coli*) and pMF1 (*M. xanthus*) | 8755 | Oxytetracycline, Ampicillin, Kanamycin |
| **16** | pMxtetR_Sp107 | pMxtetR with GH3.5.6c02 protospacer 107 inserted in BamHI+KpnI site | Plasmid interference assay | ColE1 (*E. coli*) and pMF1 (*M. xanthus*) | 8757 | Oxytetracycline, Ampicillin, Kanamycin |
| **17** | pMxtetR_Sp105 | pMxtetR with GH3.5.6c02 protospacer 105 inserted in BamHI+KpnI site | Plasmid interference assay | ColE1 (*E. coli*) and pMF1 (*M. xanthus*) | 8755 | Oxytetracycline, Ampicillin, Kanamycin |
| **18** | pMxtetR_Sp81 | pMxtetR with GH3.5.6c02 protospacer 81 inserted in BamHI+KpnI site | Plasmid interference assay | ColE1 (*E. coli*) and pMF1 (*M. xanthus*) | 8755 | Oxytetracycline, Ampicillin, Kanamycin |
| **19** | pMxtetR_Sp1 | pMxtetR with DK1622 protospacer 81 inserted in BamHI+KpnI site | Plasmid interference assay | ColE1 (*E. coli*) and pMF1 (*M. xanthus*) | 8755 | Oxytetracycline, Ampicillin, Kanamycin |

Table 3. Primers

| No. | Name | Sequence (5' to 3') | Source | Purpose |
|---|---|---|---|---|
| **1** | cas3_For | TCATCTGCTGCGCGA GCACTTGGAG | Microsynth | Amplification GH3.5.6c02 *Cas3* gene |
| **2** | cas3_Rev | CCCGGCTCGCATCGA AGAAGGCTTC | Microsynth | Amplification GH3.5.6c02 *Cas3* gene |
| **3** | GV1002 | TAATCGTCTAGAttcaatcgtcacccttttctcg | Microsynth | Amplification pMR3629 *tetR* gene |
| **4** | GV1003 | AAGTGGATCCcagatccgtccctatcgtt | Microsynth | Amplification pMR3629 *tetR* gene |
| **5** | GV1006 | gatcACAAGATCTGGCCGGTGACCCTCC GGGCTCTTGAGAAGgtac | Microsynth | GH3.5.6c02 Sp121 oligo1 |
| **6** | GV1007 | CTTCTCAAGAGCCCGGAGGGTCACCG GCCAGATCTTGT | Microsynth | GH3.5.6c02 Sp121 oligo2 |
| **7** | GV1078 | gatcCGTCGACATCGCAAGCCCGGCGA AGGACAACTGGAAGgtac | Microsynth | GH3.5.6c02 Sp120 oligo1 |
| **8** | GV1079 | CTTCCAGTTGTCCTTCGCCGGGCTTGC GATGTCGACG | Microsynth | GH3.5.6c02 Sp120 oligo2 |
| **9** | GV1080 | gatcAATTACGCTGCATCAGCACGTCCG TGCGCTAGCTGAAGgtac | Microsynth | GH3.5.6c02 Sp119 oligo1 |
| **10** | GV1081 | CTTCAGCTAGCGCACGGACGTGCTGAT GCAGCGTAATT | Microsynth | GH3.5.6c02 Sp119 oligo2 |
| **11** | GV1084 | gatcTGAATGGTGTTCATCGGATGTTGATC CCCGGCTGCTGGAAGgtac | Microsynth | GH3.5.6c02 Sp117 oligo1 |
| **12** | GV1085 | CTTCCAGCAGCCGGGGATCAACATCCG ATGAACACCATTCA | Microsynth | GH3.5.6c02 Sp117 oligo2 |
| **13** | GV1086 | gatcACCTTCACCGGGCTGATCGCGGAC AACAGCATCGTGAAGgtac | Microsynth | GH3.5.6c02 Sp115 oligo1 |
| **14** | GV1087 | CTTCACGATGCTGTTGTCCGCGATCAG CCCGGTGAAGGT | Microsynth | GH3.5.6c02 Sp115 oligo2 |
| **15** | GV1088 | gatcCGCGCGGGCCGCTTCGAGGTGGT GCGCAAGGGCGTGAAGgtac | Microsynth | GH3.5.6c02 Sp113 oligo1 |
| **16** | GV1089 | CTTCACGCCCTTGCGCACCACCTCGAA GCGGCCCGCGCG | Microsynth | GH3.5.6c02 Sp113 oligo2 |
| **17** | GV1090 | gatcTCTTGTCTTCAGCGCTACTCATGGA CGAGTCGAGAAGgtac | Microsynth | GH3.5.6c02 Sp111 oligo1 |
| **18** | GV1091 | CTTCTCGACTCGTCCATGAGTAGCGCT GAAGACAAGA | Microsynth | GH3.5.6c02 Sp111 oligo2 |
| **19** | GV1092 | gatcATGCGATGCAATTCCATGTAAGCAT CCTTCCACTGGAAGgtac | Microsynth | GH3.5.6c02 Sp110 oligo1 |
| **20** | GV1093 | CTTCCAGTGGAAGGATGCTTACATGGAA TTGCATCGCAT | Microsynth | GH3.5.6c02 Sp110 oligo2 |
| **21** | GV1040 | gatcCTCAGCGAACGTGTAGAACTCATC CACGTGGGCTCCGAAGgtac | Microsynth | GH3.5.6c02 Sp109 oligo1 |
| **22** | GV1041 | CTTCGGAGCCCACGTGGATGAGTTCTA CACGTTCGCTGAG | Microsynth | GH3.5.6c02 Sp109 oligo2 |
| **23** | GV1094 | gatcGTTCGTGGTTGTGGTGGCCAAGGC CTGCGCTGAGAAGgtac | Microsynth | GH3.5.6c02 Sp108 oligo1 |
| **24** | GV1095 | CTTCTCAGCGCAGGCCTTGGCCACCAC AACCACGAAC | Microsynth | GH3.5.6c02 Sp108 oligo2 |
| **25** | GV1096 | gatcTTCCGTGCTGTAGCGGCACTCAAC CCACTTCAAGCGAAGgtac | Microsynth | GH3.5.6c02 Sp107 oligo1 |
| **26** | GV1097 | CTTCGCTTGAAGTGGGTTGAGTGCCGCT ACAGCACGGAA | Microsynth | GH3.5.6c02 Sp107 oligo2 |

| | | | | |
|---|---|---|---|---|
| **27** | GV1098 | gatcACGCGCCAGGCCCGCGAAGGTCC GCACTGCGGGGAAGgtac | Microsynth | GH3.5.6c02 Sp105 oligo1 |
| **28** | GV1099 | CTTCCCCGCAGTGCGGACCTTCGCGG GCCTGGCGCGT | Microsynth | GH3.5.6c02 Sp105 oligo2 |
| **29** | GV1100 | gatcCTCATCCAGCACTGGGTGCATCCG AAACGTCACGAAGgtac | Microsynth | GH3.5.6c02 Sp81 oligo1 |
| **30** | GV1101 | CTTCGTGACGTTTCGGATGCACCCAGT GCTGGATGAG | Microsynth | GH3.5.6c02 Sp81 oligo2 |
| **31** | GV1046 | gatcCGCCCACCCGTCCACCACCCGGT GCGCCTCGCGGAAGgtac | Microsynth | DK1622 Sp1 oligo1 |
| **32** | GV1047 | CTTCCGCGAGGCGCACCGGGTGGTGG ACGGGTGGGCG | Microsynth | DK1622 Sp1 oligo2 |
| **33** | GV866 | cggttcctggccttttgctgg | Microsynth | Sanger sequencing spacer plasmids |
